## Supplement for "Macromolecular and electrical coupling between inner hair cells in the rodent cochlea"

**Supplementary figures**

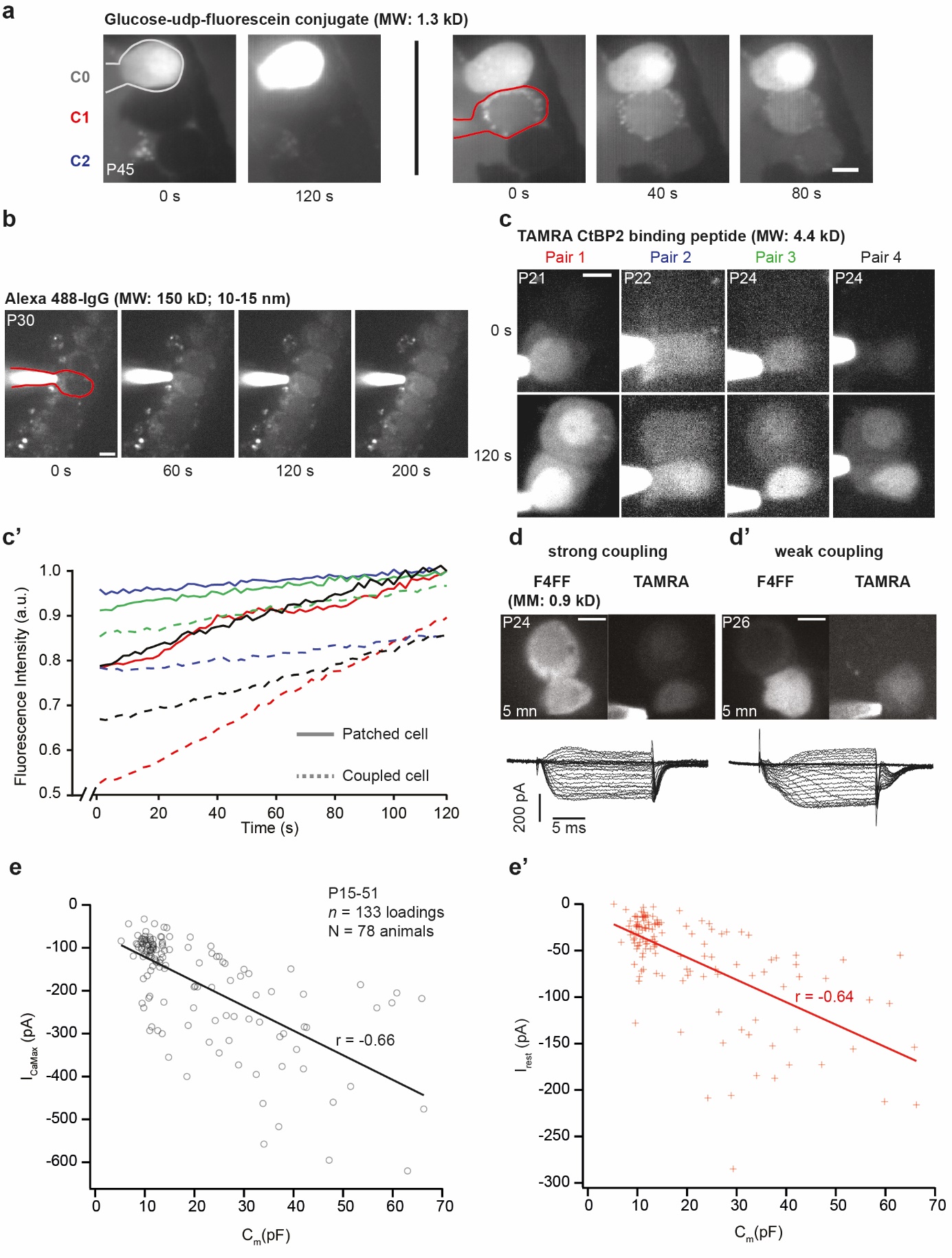

**Supplementary figure 1.** **Different levels of molecular and electrical coupling exist between IHCs.** **(a)** Time series of fluorescently labeled glucose loading in a single, non-coupled IHC (C0, no spread to other IHCs) and coupled IHC (C1) that shared glucose with the coupled IHC (C2). **(b)** Time series of the Alexa 488-IgG loading among 7 coupled IHCs. The patched IHC is delineated in red. **(c)** The pictures illustrate the TAMRA-peptide loading of four different pairs of IHCs. **(c’)** Mean fluorescence intensities of the patched IHCs (solid line) and their coupled IHCs (dashed line) over time from the loadings in (b). The fluorescence is normalized to the maximal fluorescence of the patched cell (line). **(d)** Example picture of a strong IHC-IHC coupling where the F4FF and TAMRA peptide are equally loaded in both coupled IHCs after 5 min, resulting in a monophasic activation of the whole-cell Ca^2+^-current from depolarization steps. **(d’)** The weak IHC-IHC coupling example shows much weaker intensities for F4FF and TAMRA in the non-patched cell, with a whole-cell Ca^2+^-current exhibiting multiple phases of activation. **(e-e’)** Scatter plot and line fits relating I_CaMax_ (e) and I_rest_ (e’) currents from step depolarizations against C_m_ values across all age groups from ruptured-patch experiments in 1.3,2 or 5 [Ca^2+^]_e._ *Abbreviations: a. u.,* arbitrary units. *Scale bars:* 5 µm.

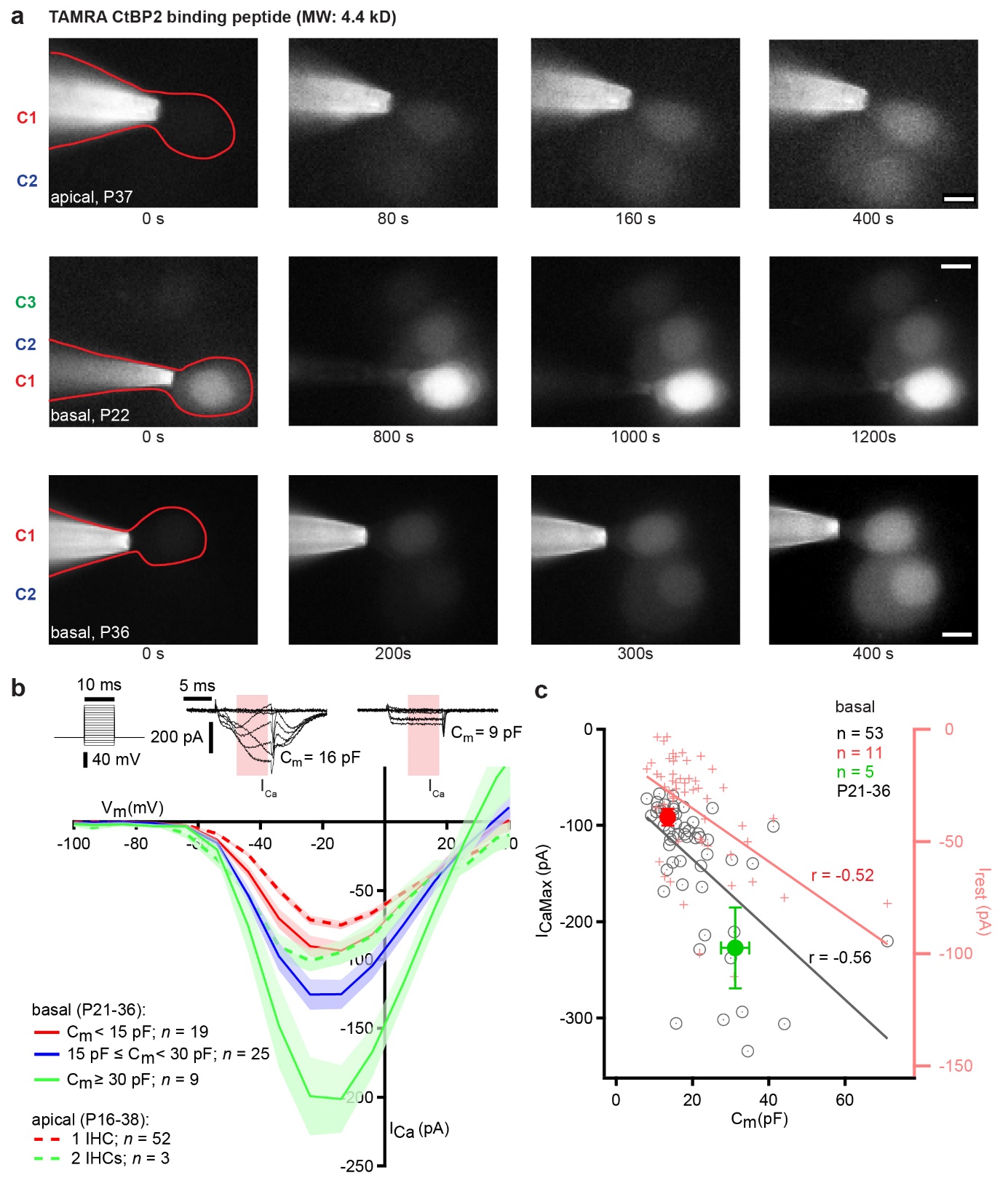
**Supplementary figure 2. Macromolecular and electrical IHC-coupling in the high- and low-frequency cochlea of the gerbil. (a)** Fluorescence images taken during loading of TAMRA-peptide through the patch pipette into the IHC. In addition to the patched cell (C1), one IHC (first row; C2) in the apical cochlea and two (second row; C2-3) IHCs or one (third row; C2) neighboring IHC in the basal cochlea were also loaded with the fluorescent dye, demonstrating a molecular coupling between these cells. Note that in the right three panels of the second row, the pipette is slightly out of focus due to drift. In the basal cochlea, the mean C_m_ of non-coupled, single IHCs (as demonstrated by the lack of TAMRA-fluorescence spread) was 13.5 ± 0.7 pF (*n* = 11 loadings). When the dye loaded into 2 or 3 neighboring cells, the C_m_ read-out was 31.0 ± 4.0 pF (*n* = 5 loadings). In the apical part the C_m_ values for non-coupled IHCs were 14.3 ± 0.3 pF (*n* = 52 loadings) and 29.8 ± 4.6 pF (*n* = 3 loadings) for coupled IHCs. Values are shown as mean ± S.E.M. **(b)** Current-voltage relationship obtained by 10-ms long depolarizations of increasing voltages (10 mV-increments) from IHCs of either basal or apical turn (N_animals_ = 42 (basal turn recordings) and N_animals_ = 17 (apical turn recordings), respectively). Ca^2+^ currents were significantly larger in dye-filled coupled cells or cells with C_m_ values from 15-30 pF and above 30 pF (likely representing coupled cells) as compared to cells with C_m_ values smaller than 15 pF. In some cells with large C_m_, the Ca^2+^ current activation displayed multiple time constants (middle insert), likely due to a delayed channel activation in the incompletely clamped neighboring cells. **(c)** The relation between the resting membrane capacitance C_m_ and the maximal depolarization-evoked Ca^2+^ current I_CaMax_ (gray circles*,* Pearson correlation coefficient: r = -0.56, p < 0.0001), as well as the resting current I_rest_ (red crosses, Pearson correlation coefficient: r = -0.52, p < 0.0001) in the basal cochlea (*n* = 53 basal cells from the panel b). Mean values for C_m_ and I_CaMax_ and their respective S.E.M. for single-dye-filled (red; *n* = 11) and coupled-dye-filled (green; *n* = 5) IHCs are presented as filled dots with error bars. Data was obtained across all age groups (P16-38) using ruptured patch experiments with 1.3 mM extracellular Ca^2+^. *Abbreviations: IHC*, inner hair cell *Scale bars:* 5 µm.

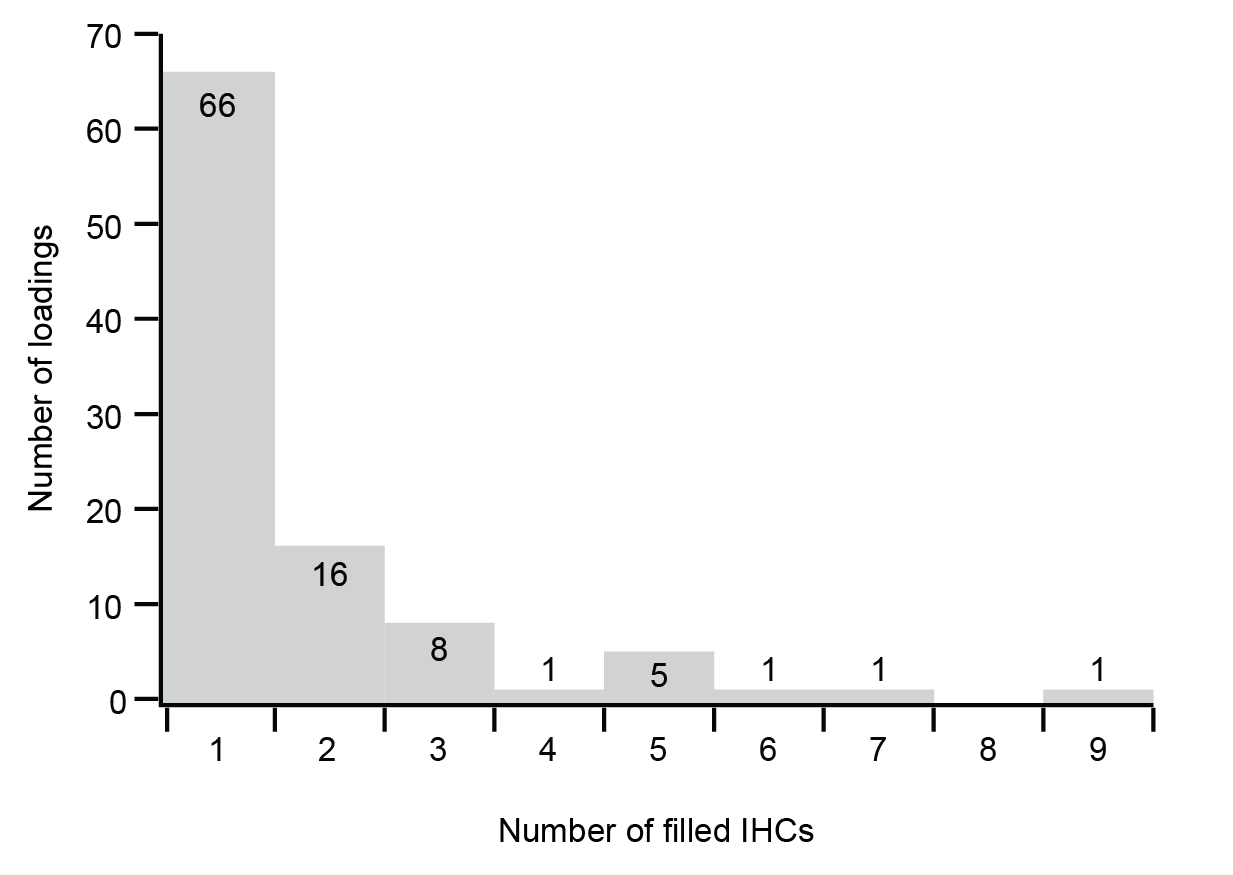

**Supplementary figure 3: Distribution of dye-filled IHC numbers in the temporal bone preparation.** Histogram displaying the number of dye-filled IHCs per loading at P26-100, with a prevalence for coupled cells of 33%. *Abbreviations: IHC*, inner hair cell.

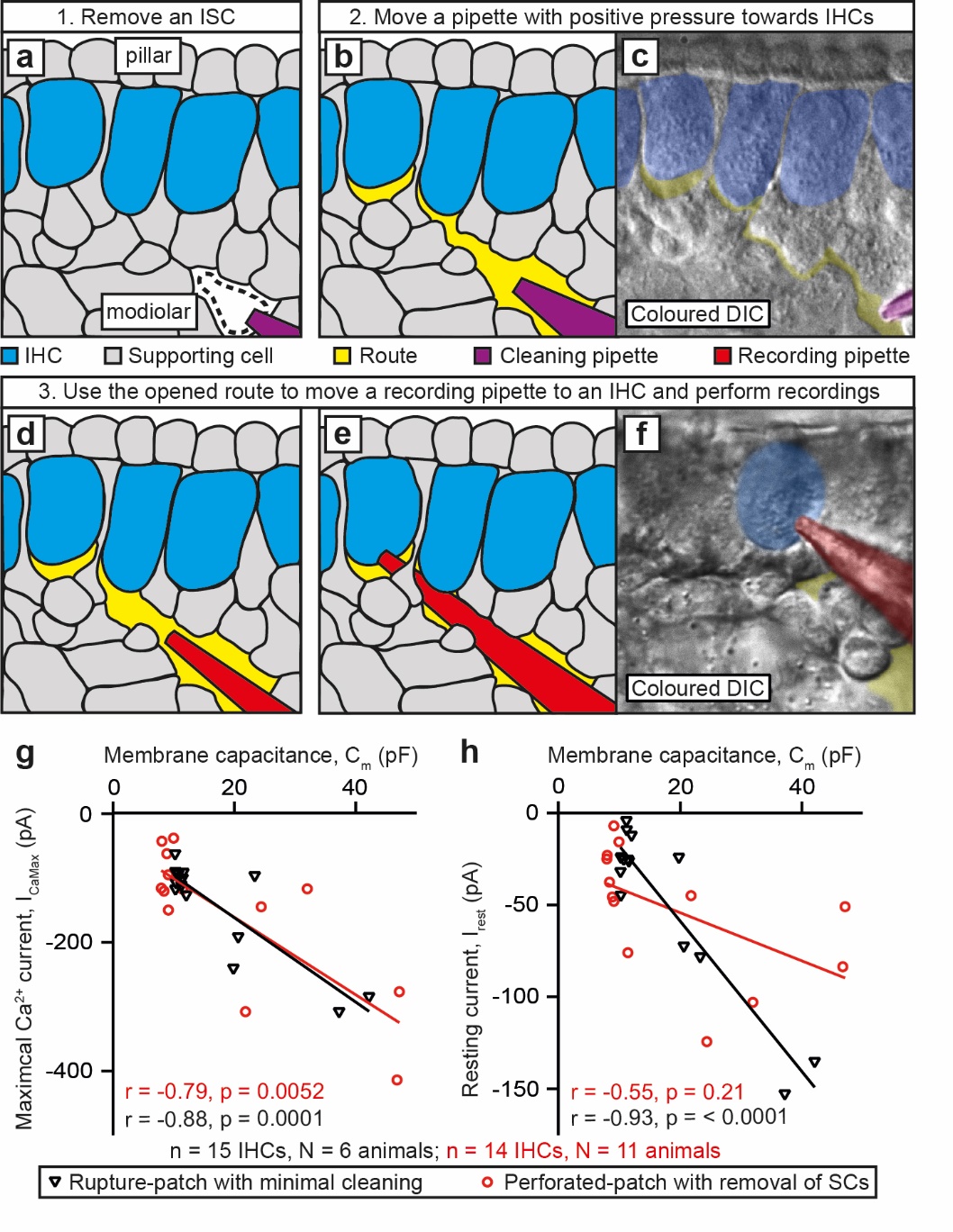

**Supplementary figure 4. IHC coupling can be observed with least invasive recording approaches.** Figures **a-f** illustrate the ruptured-patch of IHCs with minimal cleaning. (**a**) After removing a single inner sulcus cell, (**b**) a cleaning pipette with applied positive pressure is gently moved in between SCs so that a tunnel-like route that reaches IHCs is formed. (**c**) Coloured DIC image of a formed route. (**d-e**) By carefully moving a recording pipette through the route to an IHC, ruptured-patch recordings can be performed. (**f**) Coloured DIC image of a successful ruptured-patch. (**g-h**) In addition to typical IHCs with membrane capacitance (C_m_) ~ 9 pF, IHCs with atypically high Ca^2+^ currents (I_CaMax_ in g), resting membrane currents (I_rest_ in h), and C_m_ can be observed in ruptured-patch recordings with minimal cleaning (black triangle, *n* = 5 out of 15 IHCs; N_animals_ = 6, P16-P24) and with perforated-patch recordings with removal of SCs (red circle, *n* = 5 out of 14 IHCs, N_animals_ = 11, P15-P36). Linear fits are displayed (Pearson correlation coefficient for ruptured-patch with minimal cleaning: I_CaMax_: r = -0.88, p = 0.0001, I_rest_: r = -0.93, p < 0.0001; for perforated-patch with minimal cleaning: I_CaMax_: r = -0.79, p = 0.0052, I_rest_: r = -0.55, p < 0.21). *Abbreviations: DIC*, differential interference contrast, *IHC*, inner hair cell; *ISC*, inner sulcus cell; *SC*, supporting cell.

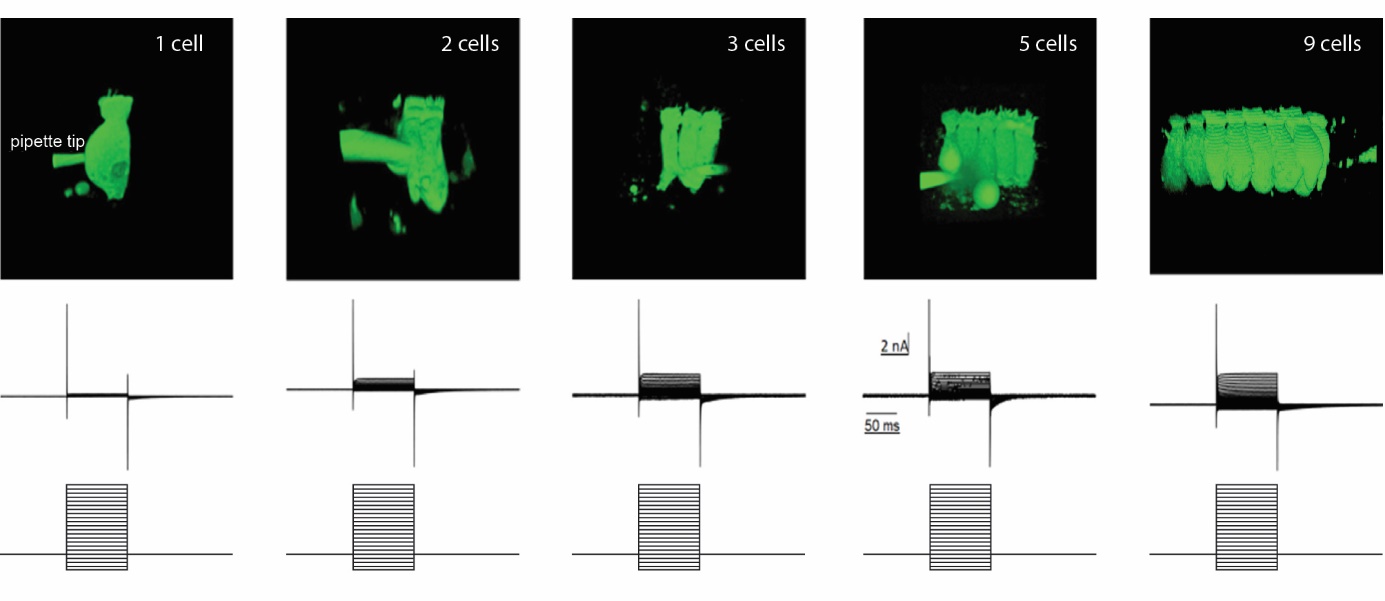

**Supplementary figure 5. Cesium outward currents increase with the number of dye-coupled IHCs.** Analyzing dye and electrical IHC coupling in temporal bone preparation, using 2PLSM imaging of OGB5N-loaded IHCs (top panel) and IHC recordings of Cs^+^-mediated outward currents (middle panel) in response to 100 ms long depolarizations (in 10 mV increments, bottom panel) from a holding potential of -60 mV. As shown for a sample of *n* = 56 distinct experiments and quantified in Fig. S6, the maximal Cs^+^ current increases with the number of dye-filled IHCs, indicating a partial summation over the currents of the coupled IHCs. We note that the recordings underestimate the true currents due to voltage drops over the series resistance to the patched IHC and the junctional resistance among the coupled cells.

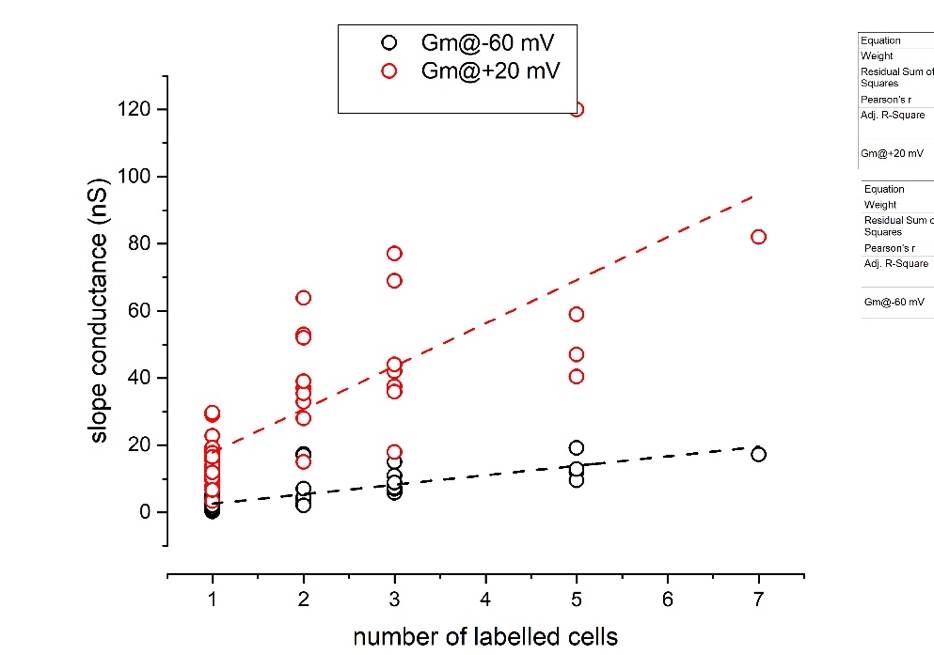

**Supplementary figure 6. IHC membrane conductances increase with the number of dye-coupled IHCs.** Membrane slope conductances (G_m_) at the holding potential -60 mV and at +20 mV from I-Vs for positively identified uncoupled and coupled cells. All the data was collected within 60 s of whole cell break-in with Cs^+^ as the major pipette cation. The membrane slope conductance at -60 mV, increasing with the number of coupled cells, is a good estimator of the effect of coupling. In this sample, using internal Cs^+^, G_m_ = 2.05 ± 1.25 nS (mean ± SD, *n* = 25 uncoupled cells) at -60 mV. Preliminary data using K^+^ as the major pipette cation did not permit the sufficient voltage-clamp control to estimate G_m_ at +20 mV. Dashed lines show regression fits, with slopes significantly different from 0 nS/#cell (Pearson correlation coefficient: r = -0.78 @ -60mV and -0.77 @ +20 mV, p<0.05).

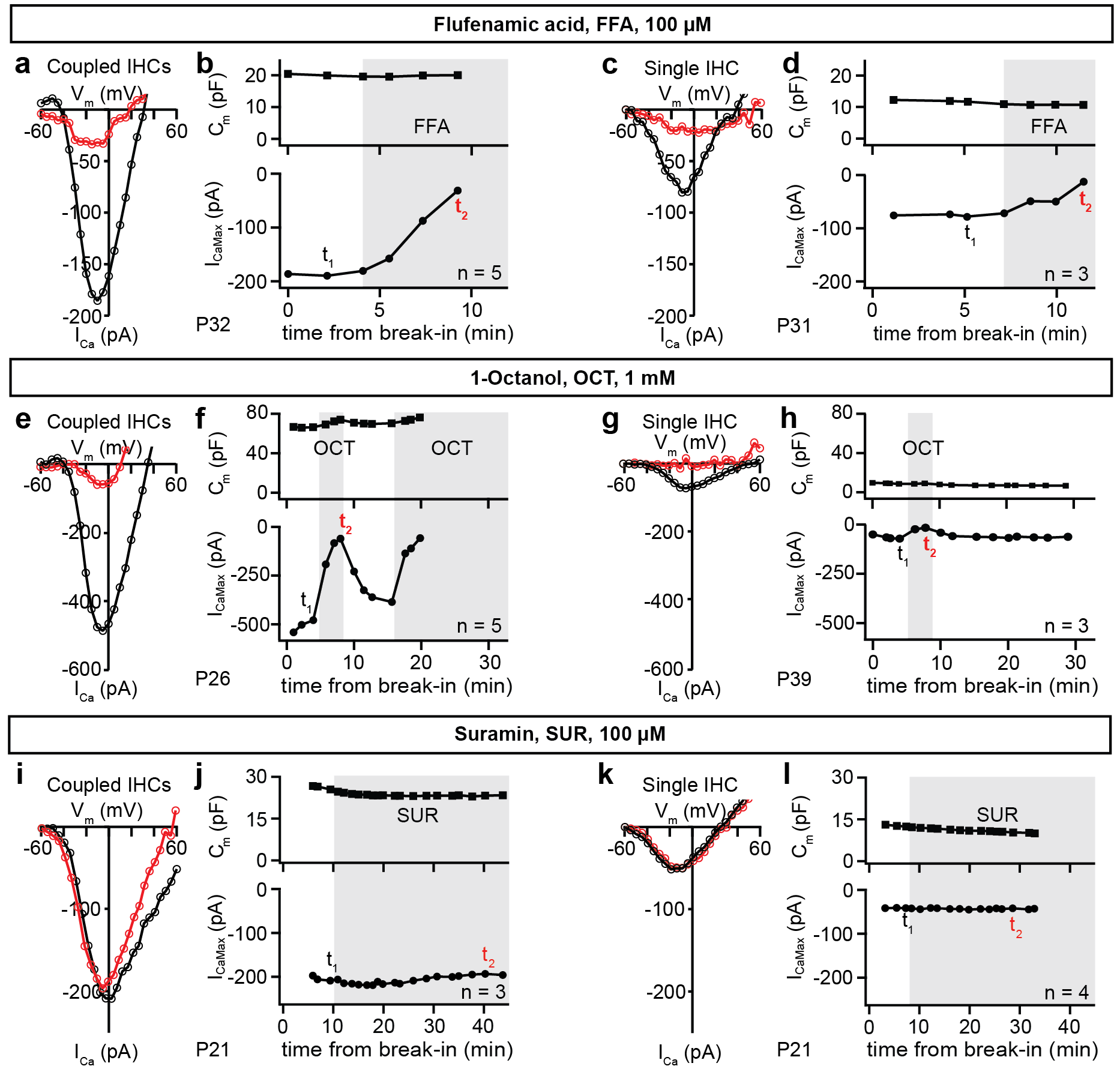

**Supplementary figure 7. Electrical IHC-coupling is not prevented by GJ blockers and suramin.** Ca^2+^-current voltage (IV) curves (a, c, e, g, I, k) were obtained by depolarizing IHCs from the holding voltage (-60 mV) with 5 mV steps before and during bath application of blockers. The presence of the blockers is marked with gray in the membrane capacitance (C_m_)-time and maximal I_Ca_ (I_CaMax_)-time plots (b, d, f, h, j, l). The recording time points of the presented IV curves are marked with t_1_ (for black curve) and t_2_ (for red curve) in the I_CaMax_-time plots. The experiments were stopped when the patched IHC was lost. **(a-h)** FFA (a, b) and OCT (e, f) inhibit I_Ca_ of coupled IHCs without changing C_m_, as seen in these representative experiments. As similar results were obtained with uncoupled IHCs (c, d, g, h), the observed reduction of I_Ca_ is not due to inhibition of IHC-coupling, but potentially due to the blockade of IHC Ca^2+^ channels. **(i-l)** As can be seen from the I_Ca_-V_m_ curves, C_m_-time, and I_CaMax_-time plots of representative experiments, suramin does not have an obvious effect on C_m_ or I_CaMax_ of coupled (i, j) and single IHCs (k, l). *Abbreviations: IHC*, inner hair cell, *FFA*, flufenamic acid; *OCT*, 1-octanol, *SUR*, suramin.

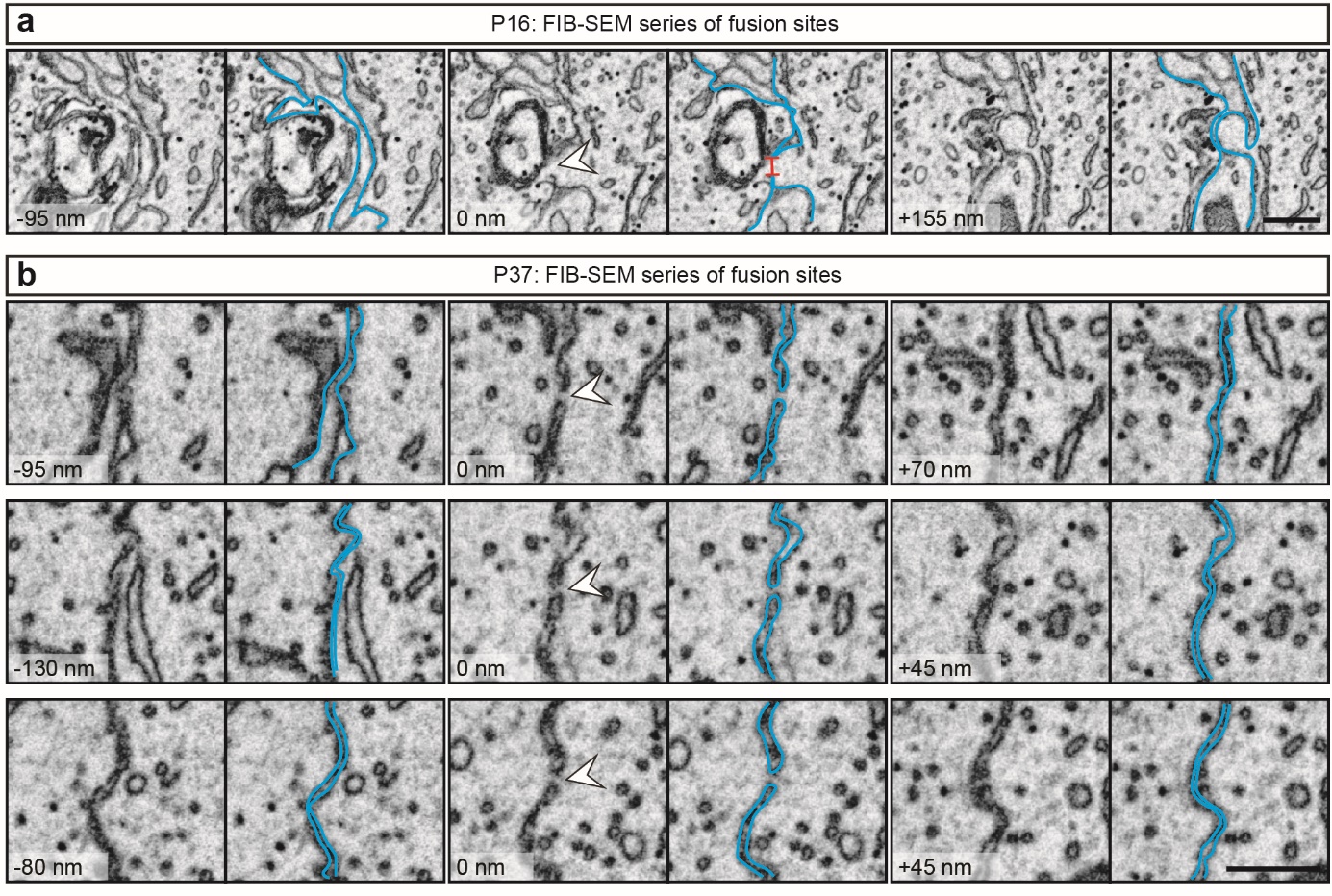
**Supplementary figure 8. FIB-SEM datasets containing putative fusion sites.** Depicted are series of putative IHC fusion sites occurring in P16 **(a)** and P37 **(b)** IHCs. Single FIB-SEM micrographs show the opposing IHC membranes (highlighted in blue) forming contacts (P16, from a filopodium; P37, from flat contacts) that contain IHC fusion sites (white arrowheads). *n* FIB-SEM run (P34/37) = 3, N_animal_ = 2; *n* FIB-SEM run (P15/16) = 3, N_animal_ = 2. For each age group 2 independend embeddings were performed. *Abbreviations: IHC*, inner hair cell. *Scale bar* = 500 nm.

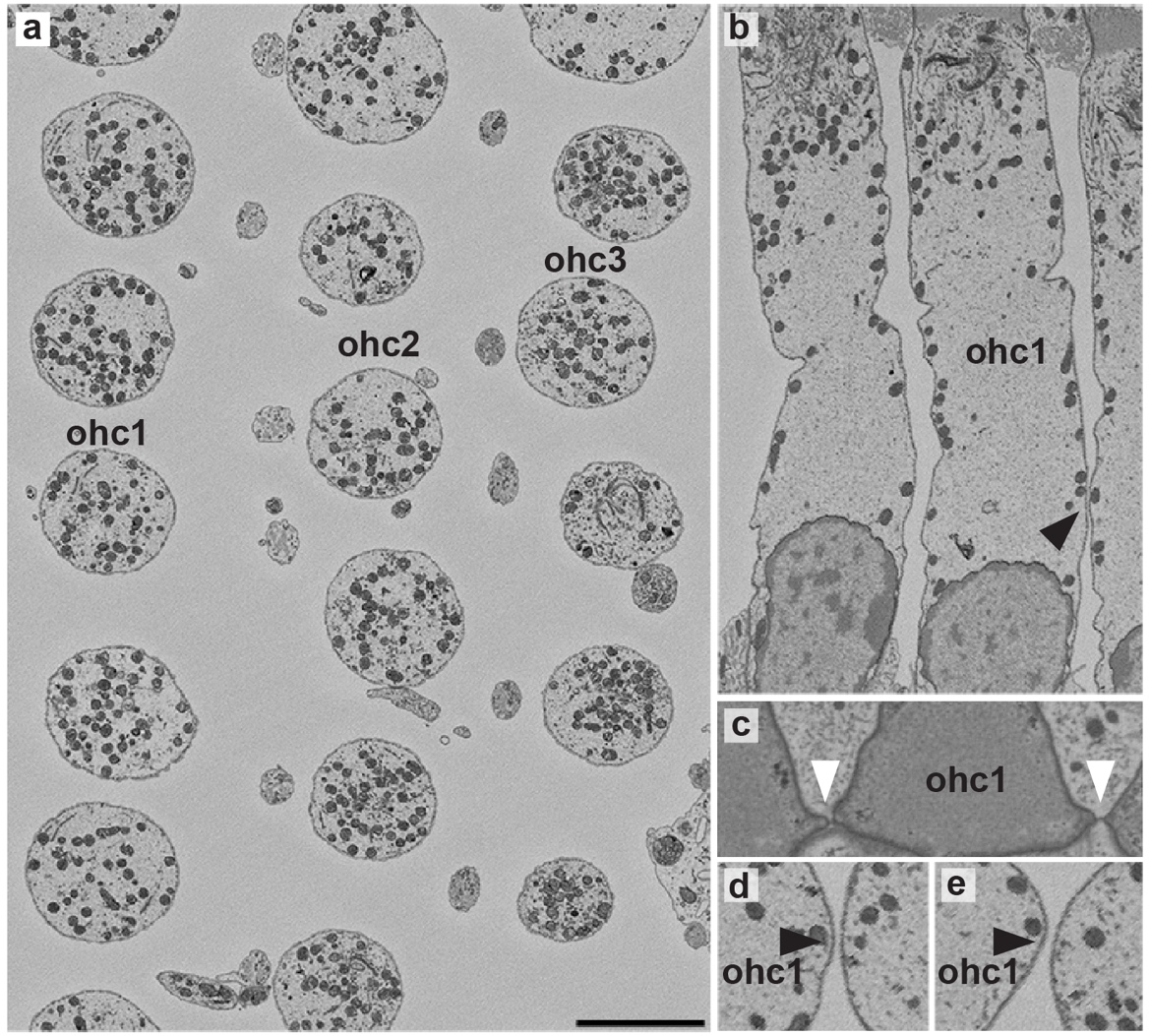

**Supplementary figure 9. No direct contacts between P22 OHCs. (a-b)** Oblique slices through a serial block-face electron microscopic dataset show only rare occasions of closely apposed basolateral membranes of OHCs (black arrowhead). **(c)** OHCs of the first row are closely apposed near their apical junctions and (**d,e**) occasionally at their basolateral membranes (black arrowheads). No direct OHC-OHC contacts were observed in two independent datasets. *Abbreviations: OHC*, outer hair cell *Scale bar* (depicted in a) = 5 µm in **a,b**; 3.3 µm in c; 2.6 µm in d,e.

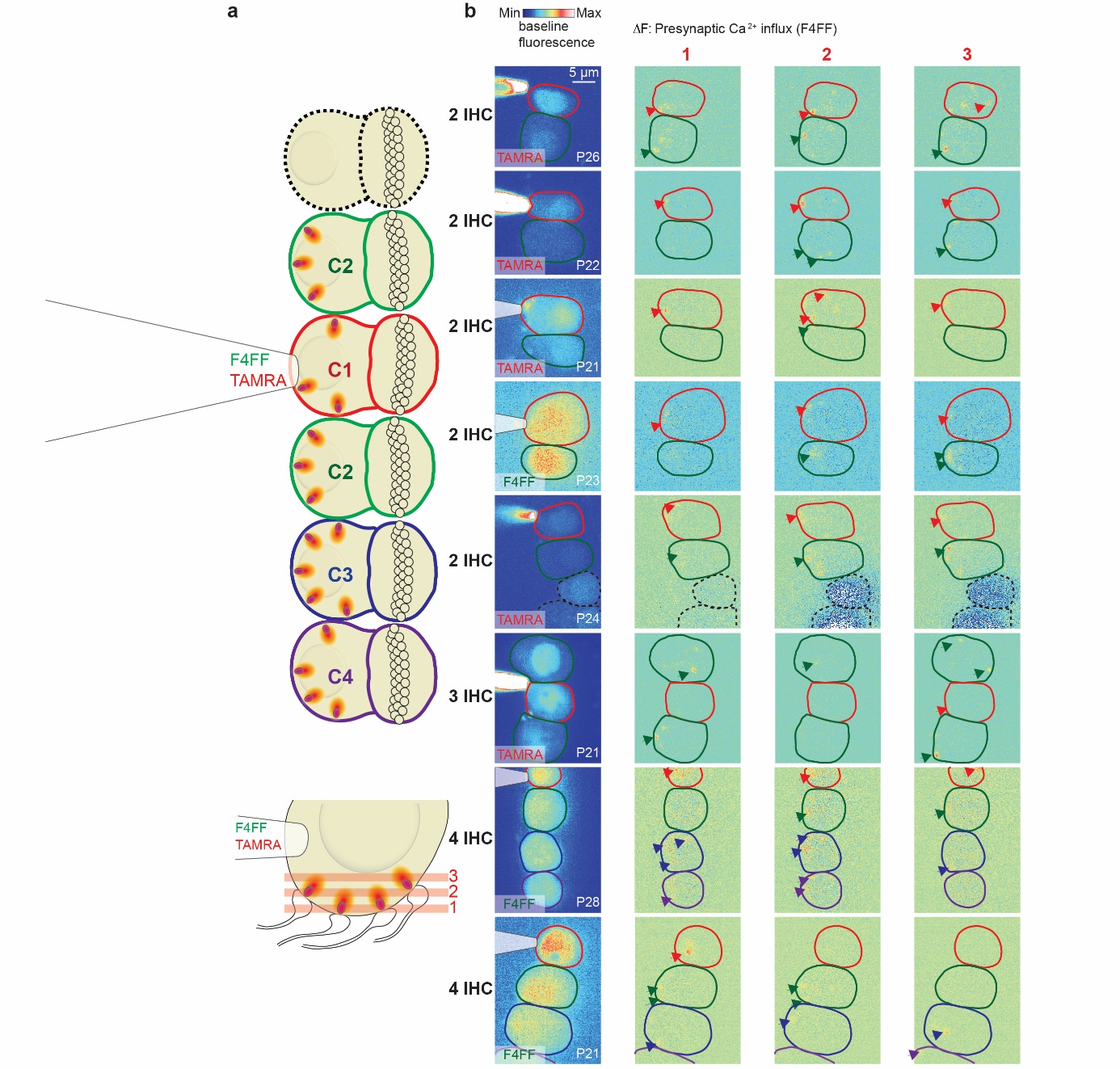

**Supplementary figure 10. Collective synaptic activity in IHC mini-syncytia.**

**a)** Coupled IHCs (delineated in color) were patch-clamped and loaded with the calcium indicator Fluo-4FF (F4FF) and the TAMRA-fluorescent CtBP2-binding peptide. C1 (red) is the patched IHC, C2 (green) are coupled IHCs neighbouring C1, C3 (blue) is coupled to C2 and C4 (violet).  Red lines show the imaging planes at bottom (1), middle (2) and top sections (3) of the synaptic IHC pole. **(b)** Left column: basal fluorescence indicating the cells in either the red (TAMRA peptide) or green (F4FF) channels. The pipette was drawn in the case it was not in the imaging plane. Right columns: ΔF/F_0_ images from single planes showing localized Ca^2+^ signals (arrowheads) in the coupled IHCs. Non-coupled IHCs that have been previously patched are delineated in dashed lines.

**Supplementary Note 1**

Encoding of auditory signals at IHC-SGN synapses entails 1) mechanoelectrical transduction at the tips of the stereocilia generating the receptor potential and 2) the coupling of the receptor potential to glutamate release (stimulus-secretion-coupling) at the ribbon synapses. Both processes involve relatively small numbers of signaling proteins: around 40 mechanotransducer channels per hair cell (*13*) and around 10 release sites per synapse (*14*). So, although neighboring IHCs receive virtually identical mechanical stimuli, because basilar membrane tuning is much wider than an IHC (see Supplementary Text 2), each IHC is subject to different realizations of stochastic mechanotransduction currents and hence they show different membrane voltage trajectories. As a consequence, signal encoding precision will be limited by the count statistics of the stochastically operating elements. The low-conductance electric coupling between the IHCs in a mini-syncytium causes voltage averaging, which increases the signal to noise ratio. This can be shown by computational modeling two scenarios in the Neuron simulation environment (*15*), three non-coupled IHCs (Fig. 8, left panel) and a mini-syncytium of three IHCs (Fig. 8, right panel), during weak stimulation driving the opening of on average 2.7 of the 40 mechanotransducer channels in an IHC (Fig. 8). Note how weakly the mechanotransducer activity is correlated and how correlations downstream of mechanotransduction are increased by IHC-coupling. Ca^2+^ channel gating seems more representative of the input in coupled than in non-coupled IHCs, demonstrated by the tendency toward higher correlation between mechanical input and Ca^2+^ channel open probability (Pearson correlation coefficient: 0.88 for coupled IHCs vs. 0.82 for non-coupled IHCs).

A simple estimation shows how coupling improves the signal to noise ratio. Assuming unmodulated weak input, the number of open mechanotransducer channels would fluctuate around an average value, e.g. four out of the 40 per IHC. At any time, the actual number of open channels is random, following a binomial distribution ℬ (N = 40, p = 0.1) with a standard deviation of *SD = sqrt (N · p · (1 - p))* = 1.90. In the extreme case of three completely fused cells, a common membrane voltage would be controlled by all 120 mechanotransduction channels. The open channel count would follow ℬ (N = 120, p = 0.1), with 12 channels open on average and an SD of 3.29. Due to the coupling, the total number of mechanotransducer channels controlling the syncytium’s membrane voltage becomes less noisy, the ratio between standard deviation and average drops from 0.47 to 0.27. The improved signal to noise ratio carries over to the fluctuations in the membrane voltage, and Ca^2+^ channel opening. Therefore, the timing of vesicle release would be more synchronized across the synapses of three coupled IHCs as compared to synapses of neighboring but non-coupled IHCs. We determined the magnitude of this effect by simulation of mechanotransduction and Ca^2+^ channel gating in non-coupled and coupled IHCs using stochastic ion channel models in the Neuron simulation environment (all code is published on the Neuron database).

**Model layout**

The simulation comprises of several IHCs, which are either isolated from each other or coupled by fusion sites. The passive properties of the IHC and the connections follow standard properties: the specific membrane capacitance, c_m_ = 10 fF/µm²; intracellular resistivity r_i_ = 120 Ohm**⋅**cm and the specific membrane resistance, r_m_ = 250 Ohm**⋅**cm² of the IHCs is chosen to obtain an effective membrane time constant of 250 µs (*16*). This basal conductance subsumes all channels that are open at rest. IHC-IHC fusion sites are approximated by 100 nm long, 78 nm wide tubes. This size is chosen to approximate a junctional resistance, R_j_ of 25 MOhm. For the numerical simulations, a cell is represented by 11 segments, a joint by 3 segments. Joints on different sides of an IHC are longitudinally offset so that they do not connect to the same segment. Stochastic gating of mechanotransducer channels and Ca_V_1.3 voltage dependent Ca^2+^ channels, are simulated with a variant of the Gillespie-Algorithm (*17*) that was adapted to work with the fixed time step of neuron simulations. To obtain statistics over many realizations of the Ca_V_1.3 gating, a two-step process is used. During the first run, the mechanotransducer channels are driven by the mechanical stimulus and gate stochastically. The number of open mechanotransducer channels is recorded at every time step. During the next thousands of runs, those open channel counts are played back while the gating of Ca_V_1.3 follows independent stochastic realizations. The output of these simulations, i.e. Ca_V_1.3 open events, are further processed to obtain release events and AP times, as described below.

**Mechanotransducer channels and voltage-gated Ca^2+^ channels**

Forty mechanotransducer channels mediate mechanosensitive influx of cations (mostly potassium) at the top end of each cell. They are driven by an external sinusoidal stimulus representing mechanical displacement in units of nanometers. A mechanotransducer channel has two states (O, C), the speed and mechano-dependence of the transition are chosen to obtain properties as described in experimental studies (*13*, *18*), i.e. sub-millisecond opening and closing transitions and an open probability that increases sigmoidally with the displacement d. The range of displacement for which the open probability changes, is over-estimated in experiments using glass probes and it is under-estimated when using the fluid jet to stimulate (*13*). Therefore, the value used here lies in between those experimental estimates. The opening and closing rates in ms^-1^ are $k_{+}=k_{max}/\left( 1+exp\left( \left( s_{1/2}^{o}-d \right)/k_{o} \right) \right)$; $k_{-}=k_{max}/\left( 1+exp\left( \left( {d-s}_{1/2}^{c} \right)/k_{c} \right) \right)$ given a displacement *d*. The parameters are $k_{max}=16.9 ms^{-1}; s_{1/2}^{o}=179 nm; s_{1/2}^{c}=-10 nm;k_{o}=50.6 nm^{-1};k_{c}=51.4 nm^{-1}$. Rates and open probability are visualized in Fig. S11 We aimed to use a realistic model that closely reproduces experimental results, even though the exact details of the mechanotransducer model are in no way crucial to the outcome of the simulation. The key aspect, the stochasticity of the voltage trajectory of individual cells, results from two main determinants: the low overall number of mechanotransducer channels and their sub-millisecond gating kinetics. Both parameters are experimentally documented in multiple studies.

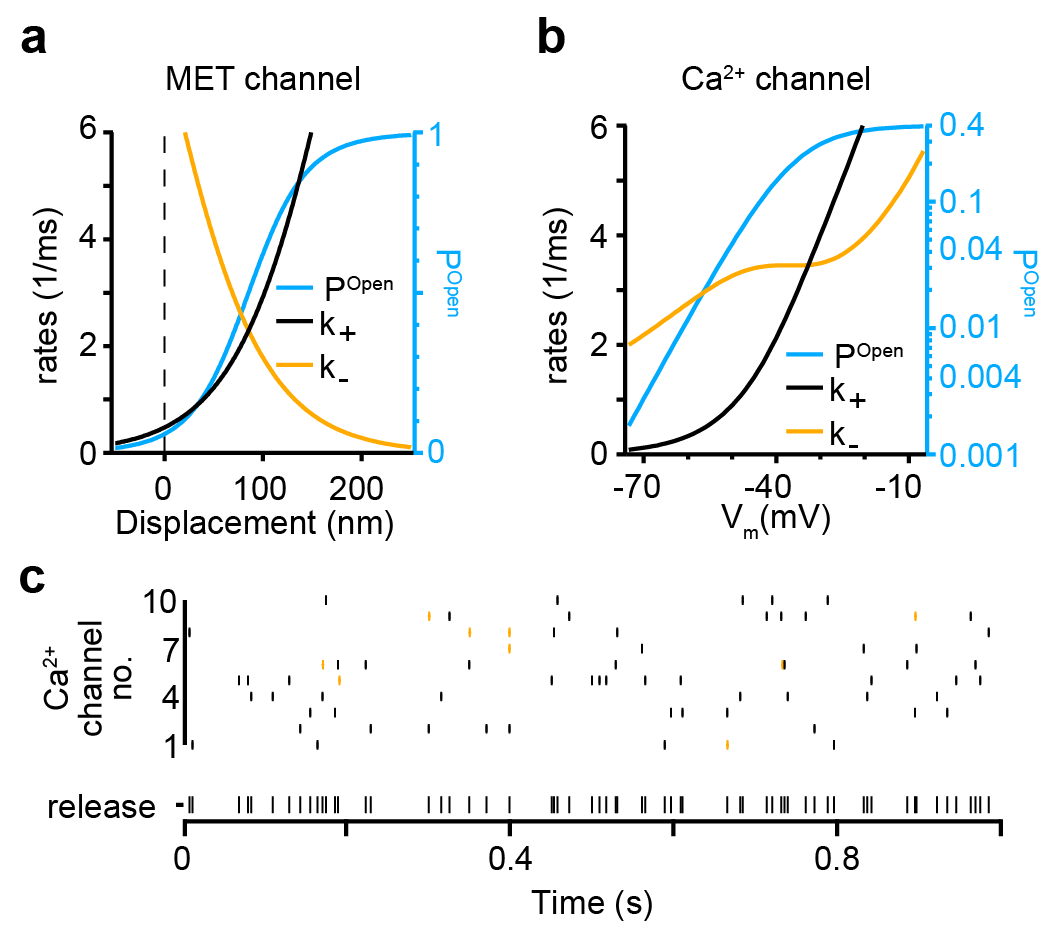

**Supplementary figure 11: Implementation of ion channels and release event pruning**. **(a)** The model for mechanotransducer channels is designed to account for published experimental result. **(b)** Ca_V_1.3 channels are implemented as described earlier (*19*), for details see text. **(c)** Approximate times of release events were determined by long Ca_V_1.3 channel openings (600 µs and longer), for which Ca^2+^ influx is expected to be sufficient to trigger release in the Ca^2+^ nanodomain-like scenario (*19*). The top ten rows show time stamps at which an opening of the respective channel (#1 to 10) exceeded a duration of 600 µs. All ten channels are from the same cell, this is, all experienced the same voltage time course. Note that only a small fraction of openings (3%) last longer than this the 600 µs cut-off. When all events from the ten channels are combined to approximate release events and then postsynaptic action potentials at the IHC ribbon synapse, nine out of 71 events are pruned (orange) when post-synaptic refractoriness is considered. The postsynaptic spike rate for this weak stimulus is on average 64 action potentials/s.

The Ca_V_1.3 Ca^2+^ channel gating parameters are obtained from experiments (*19*–*21*), derived as described before. In short, the channel has two closed and one open state with the following transitions:

$$C_{1}\begin{matrix} \underset{\to}{\text{2}k_{+}} \\ \overset{\leftarrow}{k_{-}} \end{matrix}C_{2}\begin{matrix} \underset{\to}{k_{+}} \\ \overset{\leftarrow}{{2k}_{-}} \end{matrix}O$$

with opening and closing rates in ms^-1^:

$k_{+}=\frac{\exp\left( -b\cdot V \right)}{a}\sqrt{p_{max}/\left( 1+exp\left( \left( V_{1/2}-V \right)/w \right) \right)}$ ; $k_{-}=\frac{\exp\left( -b\cdot V \right)}{a}-k_{+}$

with $a=0.2565/3^{(T-23^{\circ}C)/10K} ms$; $b= -0.0295 mV^{-1}$; $p_{max}=0.4$; $V_{1/2}=-36.2 mV$ ; $w=6.8 mV$; $T=37^{\circ}C$. Rates and open probability are visualized in Fig. S11b).

**Stimulus-secretion coupling**

Ca^2+^ enters through open Ca^2+^ channels and binds to the fusion machinery of a nearby release-competent synaptic vesicle (SV), triggering its fusion. In mature IHCs, this stimulus-secretion coupling operates mostly in the Ca^2+^ nanodomain-like regime (*19*, *22*, *23*), i.e. the opening of one or very few Ca^2+^ channels controls the fusion of a given vesicle. These processes are not explicitly modelled. Instead, we use the functional properties of Ca^2+^ nanodomain signaling and approximate the occurrence of release events with the occurrence of long Ca^2+^ channel openings. Channel open times beyond a certain cut-off time are considered effective to trigger fusion of the nearby vesicle. The exact duration of this cut-off had no influence on the outcome. This was established using cut-off values between 100 microseconds and 800 microseconds. For cut-offs beyond 800 microseconds the collection of a sufficient number of events was beyond the available processor time. The results below are derived with a cut-off of 600 microseconds.

We previously estimated the number of independent vesicle release sites per IHC ribbon synapse to be around 10, based on electron microscopy studies and quantitative analysis of release rate dynamics in response to sound stimulation (*4*, *14*). Therefore, 10 Ca^2+^-channels were considered to simulate the 10 Ca^2+^ channel-release units of a synapse. Release events occurring at any of those 10 units were considered as inputs to a single postsynaptic SGN. In mammals, most SGNs receive input from only one ribbon synapse. Release events occurring in close succession were pruned by a refractory process with an absolute refractory period of 0.6 ms and a relative refractory period of 0.65 ms as we used earlier (*24*, *25*), based on our experimental estimates (*26*). In this way, 10 independent trials of single Ca_V_1.3 channels yield one realization of an action potential sequence from a synapse (Fig. S11c. In order to obtain not only a single realization, but rather an approximation of the instantaneous rate of action potentials under the given realization of mechanotransduction, we simulated 35,000 trials equaling to 3,500 synapses (Fig. 8g).

For all simulations in this study, we restrict ourselves to weak stimuli, i.e. causing low release rates. Therefore, the release sites are rarely depleted and the dynamics of release site refilling is of little influence. Hence, to simplify the model, we ignored the time required to refill the release site. We feel this is further justified, because the improved signal to noise in the syncytium is driven by averaging of the mechano­transducer currents, i.e. events upstream of vesicle release.

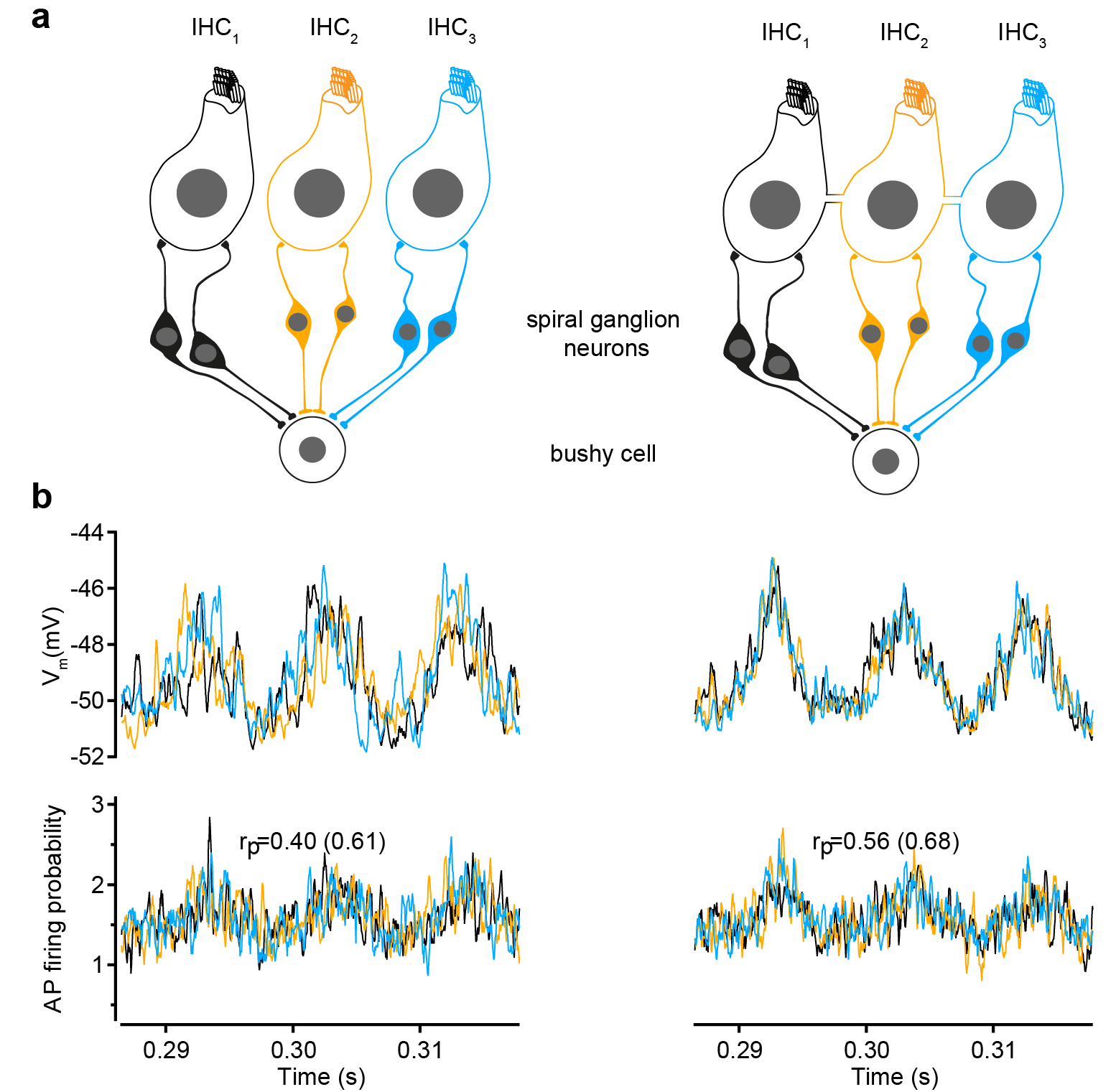

**Supplementary figure 12: Coupling of three IHCs increases the correlations between action potential firing by SGNs**. **(b)** In the first configuration of the model, six SGNs relay auditory information from three IHCs to a bushy cell, which fires only if sufficiently coincident input arrives from the SGNs. **(b)** A short section of membrane voltage and action potential frequency (average event rate within 200 µs, across 4,500 synapses) are shown for uncoupled (left) and coupled IHCs with a junctional resistance of 15 MOhm (right) - compare to the dataset for 25 MOhm in Fig. 8. Note how the coupling improves correlations between the action potential rate for SGNs originating at neighboring IHCs as well as the correlation between the mechanical stimulus and the action potential rate (in parentheses).

In summary, our model includes a deterministic sinusoidal stimulus that controls stochastic mechano­transducer channels. The resulting current drives excursions of the membrane voltage, which in turn controls Ca^2+^ channel gating and long single Ca^2+^ channel openings are considered to trigger release events. Pruned by refractoriness, release events translate to action potentials in spiral ganglion neurons. The simulation shows that coupling through 25 MOhm junctions strongly increases the correlations between the auditory input and the voltage in the coupled IHCs and also the correlations the release events (Fig. 8).

**Coincidence detection**

Although the functional benefit of coupling is clear from the increased correlation between IHC input and SGN output, we wanted to take the analysis a step further in the auditory pathway. In the cochlear nucleus multiple SGNs contact each bushy cell, which only fires, if a sufficient number of fibers is firing coincidently within a narrow time window. For example, bushy cells in the anteroventral cochlear nucleus fire an action potential, when a number of SGNs fire action potentials within a short coincidence window and, as a result, show improved precision of spike timing when compared to the spiral ganglion neurons (*27*, *28*). Each bushy cells receive multiple inputs of SGNs with similar characteristic frequency and spontaneous firing rate (*29*), which allows us to use one parameter set to simulate such a set of incoming action potential times. The number of coincidences required depends on the bushy cell type and species. It is considered to range from a few fibres up to 20 or even more. The coincidence window has no fixed width, it depends on the activity status, i.e. the depolarization history of the bushy cell (*30*). Typical bushy cell membrane time constants and spike jitter (*28*) suggest coincidence window times of 50 to 200 µs. We find that different assumptions about this coincidence window had no impact on the outcome of the model (as described below) at the cochlear nucleus (CN) because the bandwidth of signals from the IHC is already restricted by the IHC membrane time constant and the rise time of the Ca^2+^ currents.

The probability of two coincident action potentials from two SGNs within a 200 µs coincidence window simply corresponds to the product of the individual firing probabilities (Fig. S12). The average value of this product depends on the origin of the signals of the SGNs. If they received input from two synapses of the same IHC, coincidences are slightly more likely. To our knowledge, there is no information yet about the precise connectivity between IHCs and bushy cells. We studied two different configurations in which six SGNs converge onto a bushy cell, where the coincidences are detected. In the first configuration, three IHCs provide two SGN inputs each. In the second configuration, each of the six SGNs originates at a different IHC. In each case, we studied two possibilities: either all three/six IHCs are non-coupled or all three/six IHCs are coupled with each other.

For relatively weak stimuli (25 nm deflections), which one could consider close to threshold, we find typically that IHC coupling causes a subtle increase of around 2% in the coincidence rate for the three IHCs configuration and of 3% for the six IHC configuration. Interestingly, at very small mechanical stimuli, 2 nm sinusoidal deflection, when the correlation between the stimulus and the mechanotransducer signal drops below 0.06, the coupling between IHC does not cause an increase, but rather a small (1%) decrease in the coincidence rate. When the mechanotransducer channels open mostly unrelated to the stimulus, the averaging across coupled IHCs reduces the impact of the spurious channel openings on the coincidence rate. Together, the slight reduction at subthreshold stimuli and the slight increase at weak, supra-threshold inputs, serves to suppress thermal noise of the mechanotransduction by a slight increase of the detection threshold, i.e. the stimulus intensity, at which the coincidence rate crosses a certain threshold. This small effect size of a few percent, resulting from our simulations, is not unexpected given the identical mechanical input to the neighboring IHCs. It can be expected to increase, if basolateral K^+^ channels were added to the simulation of IHC conductances. These adaptation currents would provide a high-pass filter that reduces the average voltage and thereby increases the ratio between voltage modulation and signal average. The simulations presented here are kept deliberately simple to illustrate the essence of the coupling effect: a reduction in the noise contributions from the limited number of mechanotransducer channels.

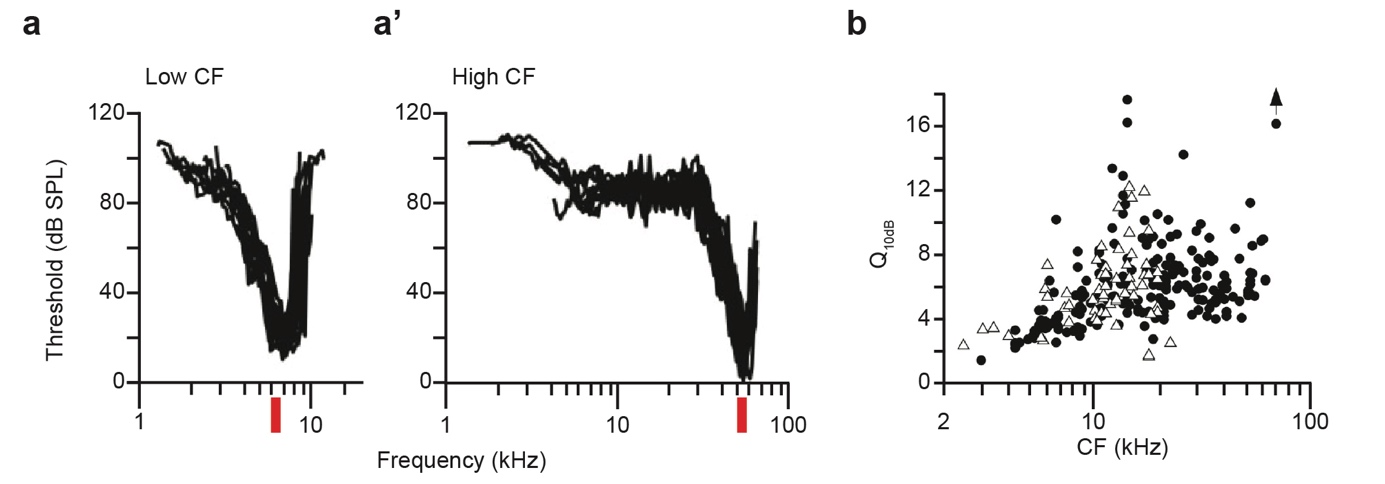
**Supplementary figure 13. Frequency resolution of sound coding by auditory nerve fibers. (a-a’)** Tuning curves of high spontaneous, low threshold auditory nerve fibers recorded from CBA/J mice taken from (*31*) and modified: **a**, auditory nerve fibers with relatively low characteristic frequency (CF) of around 7 kHz; **a’**, auditory nerve fibers with relatively high characteristic frequency (CF) of around 50-60 kHz. Red boxes indicate the frequency range covered by the largest observed mini-syncytium (9 IHCs: 83.7 µm, approximately 1.29% of the length of the organ of Corti or 0.06 octaves). **(b)** Quality of frequency tuning assessed as Q10dB (see below) of auditory nerve fibers recorded from CBA/J (circles) and C57Bl/6 (triangles) mice taken (*31*).

**Supplementary note 2**

**Does IHC-coupling degrade the frequency resolution of cochlear sound coding?**

IHC coupling with large-conductance intercellular coupling would be expected to degrade frequency resolution of sound coding if it was not limited to within the mini-syncytia that are embedded in the row of IHCs of which most (70%) are non-coupled. In our data set on P30-45 mice, the average number of cells per syncytium was 3 (3.05 ± 0.17 (S.D. = 1.53)). The mouse cochlea (with differences between strains and methods) is approximately 6-7 mm long (*32*–*34*), covers approximately 5 octaves (*35*) and contains 660-760 IHCs (*32*, *33*), each of them approximately 9 µm in diameter. Here, assuming the above parameters being 6.5 mm, 5 octaves and 700 IHCs (i.e. 9.3 µm per IHC) we calculate that the average mini-syncytium (27.9 µm) covers 0.43% of the length or 0.02 octaves. In the most extreme case observed, i.e. the largest block (9 IHCs, 83.7 µm) would cover approximately 1.29% of the length or 0.06 octaves.

Next, we compared those estimates to the physiological estimates of frequency resolution at the level of the basilar membrane and auditory nerve fibers. For the basilar membrane, the Q10 dB (characteristic frequency of basilar membrane displacement at threshold, i.e. the frequency for which the least sound pressure is required to elicit the displacement, divided by the bandwidth at 10 dB above threshold) amounts to 8.7 ± 4.3 in the high frequency range of the cochlea of CBA/J mice (50-56 kHz, (*36*)). This reflects a bandwidth at the level of the basilar membrane of approximately 5.75 kHz at 50 kHz or 0.17 octaves, approximately 3-times larger than the frequency range that is represented by the largest observed syncytium (0.06 octaves).

The quality of frequency tuning of auditory nerve fibers quantified as Q10dB (ratio of the characteristic frequency at threshold, i.e. the frequency for which the least sound pressure is required to increase the firing rate, and the frequency bandwidth of the tuning curve 10 dB above threshold) typically varies between 2 and 12 in the mouse cochlea (*26*, *31*), Fig. S13). In the characteristic frequency range from 8 kHz to 16 kHz, for which we directly demonstrated IHC-coupling in young C57Bl/6J mice, the average Q10 dB was approximately 4 (*26*, *31*) Fig. S13). For 12 kHz this reflects a bandwidth at the level of the nerve of 3000 Hz, or 0.37 octave approximately 6-times larger than represented by the broadest observed syncytium (0.06 octaves).

For a comparison to psychophysical estimates of frequency resolution, we relied on human data (*37*) which states that humans can discriminate down to 7 cents of a musical semitone (cent: percentage of a semitone). We also used a model of the human cochlea (*38*) with a cochlear length of 33 mm containing 3500 IHCs of approximately 9 µm per IHC. The mid-cochlear quarter of a turn (263-407 Hz) should then hold approximately 245 IHCs. Making a linear approximation of the frequency distribution within the mid-cochlear quarter of a turn we calculated that there are approximately 30 IHCs per a musical semitone (ST). This means each IHC “would cover” approximately 3 cents on the tonotopic map. Now, let us assume a similar situation of IHC coupling for the human cochlea as we observed experimentally in the mouse cochlea. Then, even for the case that the entire cochlea was tiled by mini-syncytia of the average size of 3 IHCs reported here, this would indicate a best possible frequency resolution of 9 cents, which is close to the lowest psychophysical estimates (7 cent, (*37*)). Therefore, we conclude that low prevalence and small size of mini-syncytia avoids trading off frequency resolution against detection sensitivity.

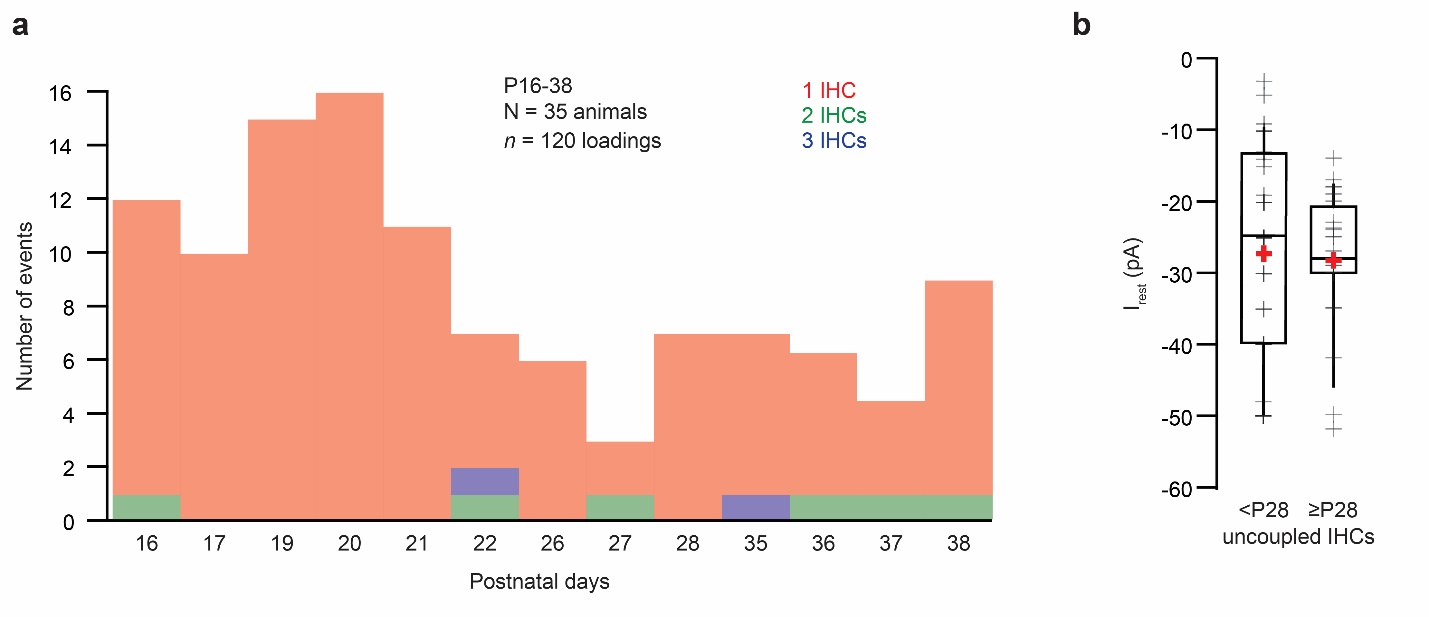

**Supplementary figure 14: Coupling probability and resting currents as a function of age in gerbils.**

**(a)** Distribution of the number of IHCs labelled during a dye-loading experiment as a function of age. **(b)** I_rest_  of un(non)-coupled IHCs before 28 and from P28 was not statistically different (<P28: -26.79 ± 2.32 pA, n  = 39, N = 15; ≥P28: -28.16 ± 1.96 pA, n  = 24, N = 9, Mann-Whitney-Wilcoxon test, p = 0.47)

| Age group | Membrane contacts/ Fusion sites | Section | Contact range in z (brackets: in xy in fusion sites) |
| --- | --- | --- | --- |
| P15  (FIB-run #1, animal 1) | 1. Flat contact IHC 1 with IHC 2  2. Filopodial contact IHC 1 with IHC 2  3. Filopodial contact IHC 1 with IHC 2  4. Filopodial contact IHC 1 with IHC 2  5. Filopodial contact IHC 1 with IHC 2  6. Filopodial contact IHC 1 with IHC 2 | 571-580  929-934  1778-1796  1781-1792  1955-1994  1973-1981 | 45 nm  25 nm  90 nm  55 nm  195 nm  40 nm |
|  | 1. Flat contact IHC 2 with IHC 3  2. Filopodial contact IHC 2 with IHC 3  3. Filopodial contact IHC 2 with IHC 3  4. Filopodial contact IHC 2 with IHC 3 | 1549-1556  1620-1726  1813-1846  2024-2028 | 35 nm  530 nm  165 nm  20 nm |
| P15  (FIB-run #2,  animal 1) | 1. Filopodial contact IHC 1 with IHC 2  2. Filopodial contact IHC 1 with IHC 2  3. Filopodial contact IHC 1 with IHC 2  4. Filopodial contact IHC 1 with IHC 2  5. Filopodial contact IHC 1 with IHC 2  6. Filopodial contact IHC 1 with IHC 2  7. Filopodial contact IHC 1 with IHC 2  8. Filopodial contact IHC 1 with IHC 2  9. Filopodial contact IHC 1 with IHC 2  10. Filopodial contact IHC 1 with IHC 2 | 3341-3360  3569-3581  3412-3454  3418-3457  3453-3459  3569-3580  3666-3677  3672-3705  3730-3747  3805-3812 | 95 nm  60 nm  210 nm  195 nm  30 nm  55 nm  55 nm  165 nm  85 nm  35 nm |
|  | 1. Filopodial contact IHC 2 with IHC 3  2. Filopodial contact IHC 2 with IHC 3  3. Filopodial contact IHC 2 with IHC 3  4. Filopodial contact IHC 2 with IHC 3  5. Filopodial contact IHC 2 with IHC 3  6. Filopodial Contact IHC 2 with IHC 3  7. Filopodial contact IHC 2 with IHC 3  8. Filopodial contact IHC 2 with IHC 3  9. Filopodial contact IHC 2 with IHC 3  10. Filopodial contact IHC 2 with IHC 3 | 1073-1088  1263-1285  1348-1375  1383-1407  1906-1928  1933-1938  2060-2148  2300-2328  2616-2656  2688-2696 | 75 nm  110 nm  135 nm  120 nm  110 nm  25 nm  88 nm  140 nm  200 nm  40 nm |
| P16  (FIB-run #3,  animal 2) | 1. Flat contact IHC 2 with IHC 3  2. Fusion site between IHC 2 and IHC 3  3. Filopodial contact IHC 2 with IHC 3  4. Filopodial contact IHC 2 with IHC 3  5. Filopodial contact IHC 2 with IHC 3  6. Filopodial contact IHC 2 with IHC 3 | 17-64  738-745  1110-1135  1245-1260  2775-2785  2820-2870 | 47 nm  35 nm (xy: 135.83 nm)  125 nm  75 nm  50 nm  250 nm |
| P34  (FIB-run #1,  animal 1) | 1. Flat contact IHC 1 with IHC 2  2. Flat contact IHC 1 with IHC 2 | 1357-1558  1615-1726 | 1 µm  555 nm |
|  | 1. Flat contact IHC 2 with IHC 3  2. Flat contact IHC 2 with IHC 3  3. Flat contact IHC 2 with IHC 3  4. Flat contact IHC 2 with IHC 3 | 11-76  453-1687  586-634  2129-2200 | 325 nm  6.17 µm  240 nm  355 nm |
| P34  (FIB-run #2,  animal 1) | 1. Flat contact IHC 1 with IHC 2  2. Flat contact IHC 1 with IHC 2 | 1-766  1144-1411 | 3.83 µm  1.335 µm |

| Age group | Membrane contacts/ Fusion sites | Section | Contact range in z (brackets: in xy in fusion sites) |
| --- | --- | --- | --- |
| P34  (FIB-run #2,  animal 1) | 3. Fusion site between IHC 1 and IHC 2 | 1233-1340 | 535 nm (xy: 124.81 nm) |
|  | 1. Flat contact IHC 2 with IHC 3  2. Fusion site between IHC 2 and IHC 3  3. Fusion site between IHC 2 and IHC 3  4. Fusion site between IHC 2 and IHC 3  5. Filopodial contact IHC 2 with IHC 3 | 1-1112  115-121  519-526  1171  1730-1768 | 5.56 µm  30 nm ( xy: 101.98 nm)  35 nm ( xy: 146.82 nm)  5 nm ( xy: 123.69 nm)  190 nm |
| P37  (FIB-run #3,  animal 2) | 1. Flat contact IHC 1 with IHC 2 | 11-22 | 55 nm |
|  | 1. Flat contact IHC 2 with IHC 3  2. Filopodial contact IHC 2 with IHC 3  3. Flat contact IHC 2 with IHC 3 | 12-25  234-275  1717-1862 | 65 nm  205 nm  725 nm |
|  | 1. Flat contact IHC 3 with IHC 4  2. Flat contact IHC 3 with IHC 4  3. Fusion site between IHC 3 and IHC 4  4. Fusion site between IHC 3 and IHC 4  5. Fusion site between IHC 3 and IHC 4  6. Fusion site between IHC 3 and IHC 4 | 304-314  1445-2128  1653-1661  1706-1709  1821-1824  2233-2244 | 50 nm  3.42 µm  40 nm (xy: 30.15 nm)  15 nm (xy: 14.42 nm)  15 nm (xy: 15.23 nm)  55 nm (xy: 52.80 nm) |

**Table S1. Quantification of contact sites and putative IHC fusion sites in the FIB data.** IHC-IHC membrane contacts are categorized into contact sites via a filopodium touching the neighboring IHC (filopodial contact) and flat membrane contacts. Both contact categories can contain perforations, likely due to the fusion of membranes of two neighboring IHCs (putative IHC fusion sites). In P34/37 IHCs, the flat contacts as well as putative IHC fusion sites are more prevalent than at P15/16 IHCs, when the filopodial contacts prevail (Table 1 for summary). The length of the individual contact sites in z is calculated via the number of sections, each section accounts for 5 nm. The xy length of the syncytial contacts was measured in the section with the largest extend of the perforation in 3dmod. In each dataset, one full IHC was visible in the region of interest, flanked by neighboring IHCs that were only partially visible, but showing the full membrane contact (see also Fig. 5). *n* FIB-SEM run (P34/37) = 3, N_animals_ = 2; *n* FIB-SEM run (P15/16) = 3, N_animals_ = 2. In total, 3 full IHCs with their contacts to neighboring IHCs were analysed per age group. Per age group 2 independent embeddings were performed. Note that 2 FIB-runs from P15 and 1 FIB-run from P34 do not show putative fusion sites.
